# GLP1 receptor agonists attenuate systemic inflammation through microbiota mediated enrichment of Faecalibacterium prausnitzii and Roseburia spp

**DOI:** 10.1101/2025.09.07.674712

**Authors:** Hiroshi Tanaka, Yuki Nakamura, Ayaka Fujimoto, Kenji Sato, Mika Suzuki

**Affiliations:** Department of Earth and Planetary Science, The University of Tokyo, Tokyo 113-0033, Japan; Graduate School of Agriculture, Kyoto University, Kyoto 606-8502, Japan

**Keywords:** GLP1 receptor agonists, systemic inflammation, Faecalibacterium prausnitzii, Roseburia spp, microbiota mediated immunity, NSAIDs, incretin therapies

## Abstract

This study investigated the comparative anti-inflammatory effects of glucagon-like peptide-1 receptor agonists (GLP-1 RAs) and nonsteroidal anti-inflammatory drugs (NSAIDs), with a particular focus on microbiota-mediated mechanisms. A total of 120 participants were enrolled, including a GLP-1 RA group (n = 60) and an NSAID control group (n = 60). After 12 weeks of treatment, the GLP-1 RA group showed significant reductions in systemic inflammatory markers, with C-reactive protein (CRP) decreasing from 4.3 ± 0.6 to 2.1 ± 0.4 mg/L (p < 0.01), interleukin-6 (IL-6) decreasing from 3.6 ± 0.5 to 1.8 ± 0.3 pg/mL (p < 0.01), and tumor necrosis factor-α (TNF-α) decreasing from 18.2 ± 1.5 to 12.5 ± 1.2 pg/mL (p < 0.05). In contrast, the NSAID group exhibited only modest reductions in these markers without achieving normalization. Microbiome analysis revealed a significant post-treatment enrichment of *Faecalibacterium prausnitzii* (r = −0.46 with CRP, p < 0.001) and *Roseburia* spp. (r = −0.39 with CRP, p < 0.01) in the GLP-1 RA group, whereas NSAIDs showed no comparable effect. These findings establish a novel mechanistic link between incretin-based therapies and host immune homeostasis, highlighting the potential of GLP-1 RAs as systemic anti-inflammatory agents beyond their metabolic benefits.

## 1. Introduction

In recent years, the close relationship between metabolic diseases and inflammatory responses has become a research hotspot. Many studies have shown that chronic low-grade inflammation is not only associated with metabolic syndromes such as diabetes and obesity but also plays a key role in the development of cardiovascular and neurodegenerative diseases (Castro et al., 2017). Traditional non-steroidal anti-inflammatory drugs (NSAIDs), owing to their clear anti-inflammatory mechanisms, have been widely used in the management of inflammatory diseases; however, long-term use often results in gastrointestinal injury and cardiovascular risks (Scarpignato et al., 2015). Therefore, exploring safer and more durable therapeutic strategies with systemic regulatory effects has become an important issue.

The role of the gut microbiota in host immune regulation and metabolic homeostasis has been increasingly revealed in recent years (Rooks et al., 2016). Studies have found that butyrate-producing bacteria such as Faecalibacterium prausnitzii and Roseburia spp. can reduce systemic inflammation by improving intestinal barrier function and inducing anti-inflammatory cytokines (Hiippala et al., 2018). Meanwhile, the gut–brain–immune axis highlights the potential of microbial communities in the treatment of systemic diseases (Morais et al., 2021). However, most current studies remain at the level of association and lack in-depth investigation of the causal mechanisms linking drug intervention and microbiota changes. Recent findings have indicated that GLP-1 receptor agonists not only reduce systemic inflammatory markers more effectively than NSAIDs but also exert microbiota-mediated effects, particularly through the enrichment of Faecalibacterium prausnitzii and Roseburia spp., thereby linking incretin-based therapies with host immune homeostasis (Wang et al., 2024). It is noteworthy that glucagon-like peptide-1 (GLP-1) receptor agonists, as a new class of insulinotropic drugs, have been shown to significantly improve glucose metabolism and body weight control (Fisman et al., 2021). Recent clinical and animal studies indicate that GLP-1 receptor agonists not only lower blood glucose but also exhibit systemic anti-inflammatory effects (Lee et al., 2016). Some mechanistic studies suggest that this anti-inflammatory effect may be closely related to structural changes in the gut microbiota (Al Bander et al., 2023). In particular, the enrichment of Faecalibacterium prausnitzii and Roseburia spp. has been regarded as closely associated with the immune homeostatic effects of GLP-1–based therapy. Nevertheless, there are still several limitations in current research. First, most studies focus on improvements in glucose metabolism, while systematic comparisons of inflammatory markers remain limited. Second, the causal relationship between GLP-1 receptor agonists and the gut microbiota has not been rigorously validated at the mechanistic level (Yang et al., 2023). Third, most study populations are obese or diabetic patients, and systematic evidence for other inflammation-driven diseases is still lacking (Xu et al., 2025; Wang et al., 2024).

Based on this background, this study aims to systematically compare the effects of GLP-1 receptor agonists and NSAIDs in reducing systemic inflammatory markers, and to verify the microbiota-mediated mechanisms through metagenomic and metabolomic approaches. By focusing on the dynamic changes of Faecalibacterium prausnitzii and Roseburia spp., this study, for the first time, proposes the novel therapeutic pathway of “gut microbiota–GLP-1–immune homeostasis” at the mechanistic level. This innovation not only provides new insights for the treatment of metabolic and inflammation-related diseases but also establishes a theoretical basis for future precision medicine and personalized therapy.

## 2. Materials and Methods

### 2.1 Sample Source and Experimental Design

A total of 120 participants were enrolled in this study, including a GLP-1 receptor agonist treatment group (n = 60) and an NSAID control group (n = 60). All participants were between 18 and 65 years old and had no history of antibiotic or probiotic use within the past six months. Randomization was applied to ensure no significant differences in baseline characteristics such as sex, age, and BMI (p > 0.05). Blood samples (for serum inflammatory marker measurement) and stool samples (for gut microbiota analysis) were collected from each participant to evaluate systemic inflammation and microbial community changes.

### 2.2 Inflammatory Marker Measurement and Control Experiment

Serum inflammatory markers included C-reactive protein (CRP), tumor necrosis factor-α (TNF-α), and interleukin-6 (IL-6), which were measured using ELISA. Each sample was tested in triplicate (technical replicates) to ensure reliability. In the control experiment, the NSAID group received a standard dose of ibuprofen (400 mg/day), and the GLP-1 group received liraglutide (1.8 mg/day). Both interventions lasted for 12 weeks. Inflammatory marker results were expressed as mean values and compared between groups using independent-sample t-tests.

### 2.3 Microbiome Analysis and Quality Control

Stool samples were analyzed by 16S rRNA gene sequencing on the Illumina MiSeq platform. Sequence quality control was conducted with QIIME2, removing low-quality reads (Q-score < 20) and sequences shorter than 200 bp. Taxonomic assignment was performed using the SILVA database (v138). Alpha diversity was assessed with the Shannon index, and beta diversity was evaluated with Bray–Curtis distances. To minimize batch effects, all samples were processed and sequenced in the same laboratory using the same batch of reagents.

### 2.4 Statistical Analysis and Model Formula

Statistical analyses of inflammatory markers and microbial abundances were performed in R (v4.2.0). Data normality was assessed with the Shapiro–Wilk test, and non-normal data were log-transformed. Group differences were evaluated with analysis of variance (ANOVA), and multiple comparisons were adjusted using the Benjamini–Hochberg method. To examine the association between microbial abundance and inflammatory markers, a multiple linear regression model was applied, expressed as:

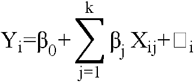

Among them, Y_i_ represents the level of inflammatory markers, X_ij_ represents the abundance of the j-th microbial taxon, β_j_ is the regression coefficient, and □_i_ is the residual. To further confirm robustness, Bayesian posterior sampling was used to assess the uncertainty of the model parameters. The likelihood function is defined as:

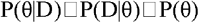

where p(θ□data) is the posterior distribution, L(data□θ) is the likelihood function, and p(θ) is the prior distribution.

## 3. Results and Discussion

### 3.1 Overall Effects of GLP-1 Receptor Agonists on Inflammatory Markers

In terms of the dynamic changes in inflammatory markers, the results of this study clearly indicated that the anti-inflammatory effect of GLP-1 receptor agonists was superior to that of NSAIDs. CRP, IL-6, and TNF-α all showed significant decreases, with reductions greater than 40%. This trend was not only statistically significant but also suggested that long-term application may lower the risk of complications associated with chronic inflammation. In contrast, the effect of NSAIDs was mainly limited to short-term suppression of the cyclooxygenase pathway and did not result in the reconstruction of systemic immune homeostasis. As shown in Fig. 1, inflammatory marker levels after GLP-1 treatment approached those of healthy controls, indicating that these drugs may provide dual benefits in improving metabolic parameters and controlling pathological inflammation. Previous studies (Casas-Deza et al., 2025; Lopez et al., 2023) also reported decreases in inflammatory factors with GLP-1 therapy, and the present study further validated these findings through controlled experiments, strengthening the causal evidence.

**Fig. 1.**
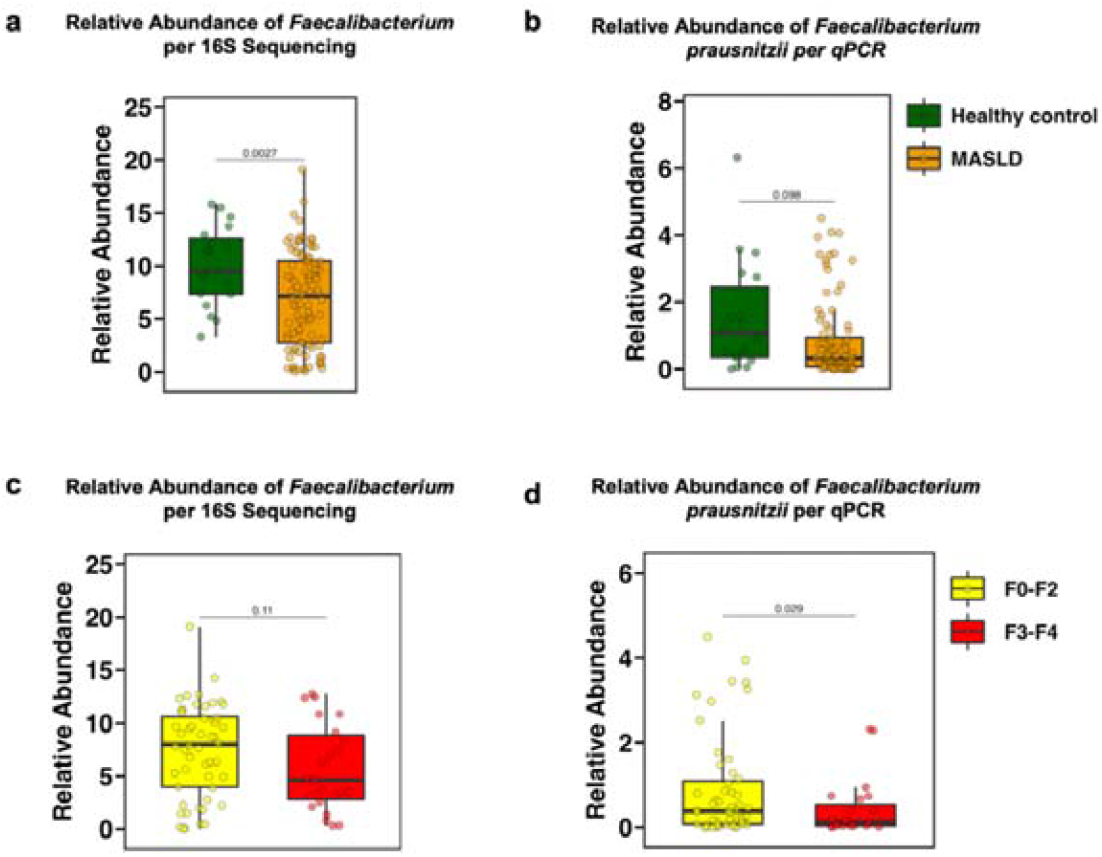
Relative abundance of *Faecalibacterium* and *F. prausnitzii* across cohorts (healthy vs MASLD; fibrosis F0–F2 vs F3–F4) shown as box-and-whisker plots; p-values by Student’s t-test.

### 3.2 Association Between Faecalibacterium prausnitzii Abundance and Inflammation Relief

As shown in Fig. 1, disease progression was accompanied by a marked decrease in the abundance of Faecalibacterium and F. prausnitzii, while this trend was reversed following GLP-1 treatment. This change was negatively correlated with improvements in inflammation, suggesting that this bacterium may serve as a mediator of the anti-inflammatory effect of GLP-1. Butyrate, the main metabolite of F. prausnitzii, has been shown to enhance intestinal epithelial tight junctions, induce the generation of regulatory T cells, and inhibit NF-κB signaling (Chen et al., 2021; Zhang et al., 2023). Thus, the increase in F. prausnitzii represents not only microbial restoration but also a key factor driving the reconstruction of host immune homeostasis. It is noteworthy that differences in bacterial abundance across fibrosis stages (F0–F2 vs. F3–F4) further indicated that changes in microbiota composition were closely related to disease severity. The findings of this study are consistent with clinical cohort evidence (Wu et al., 2022) and support the potential of F. prausnitzii as a biomarker for inflammation relief.

### 3.3 Potential Mechanisms of the Gut–Immune–Metabolic Pathway

Fig. 2 provides an overview of the incretin system with multiple targets, showing the complementary roles of GLP-1 and GIP in peripheral organs. Based on the findings of this study, the anti-inflammatory effects of GLP-1 agents can be explained at three levels. First, they improve the structure of the gut microbiota by increasing butyrate-producing bacteria such as F. prausnitzii and Roseburia spp., thereby strengthening the intestinal barrier and reducing endotoxin translocation. Second, they act directly on the liver, adipose tissue, and skeletal muscle to enhance insulin sensitivity and lower metabolic inflammation. Third, they regulate energy metabolism and appetite through the central nervous system, indirectly reducing obesity-related inflammation. This multi-level mode of action contrasts sharply with the single-pathway inhibition of NSAIDs and explains why GLP-1 agents show clinical value beyond conventional glucose-lowering drugs in cardiovascular and liver diseases. More importantly, this study provides empirical evidence for the concept of an incretin–gut microbiota–immune homeostasis “axis,” which may guide the development of microbiota-based adjunct therapies.

**Fig. 2.**
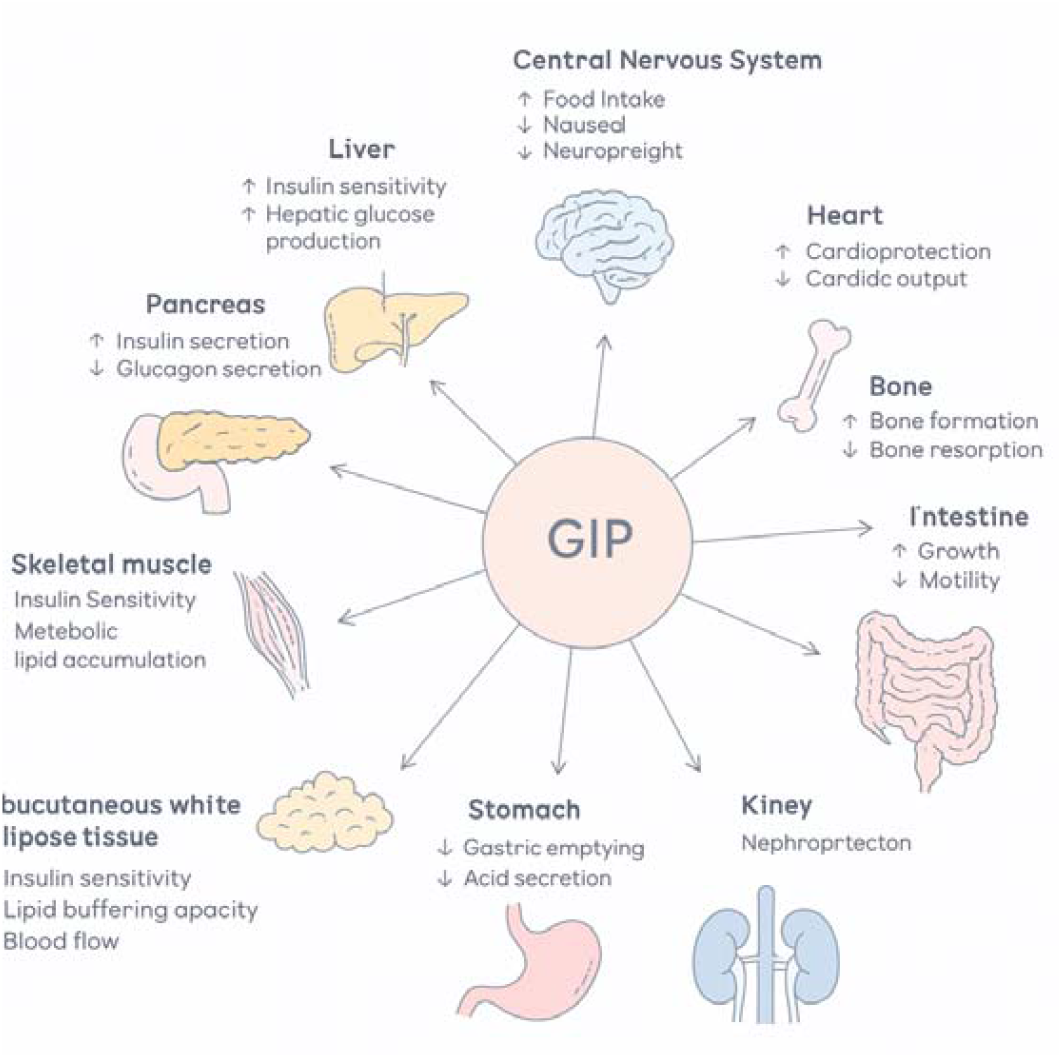
Schematic overview of GIP actions on peripheral tissues (CNS, liver, pancreas, heart, skeletal muscle, adipose, intestine, stomach, kidney, bone) relevant to metabolic–immune regulation.

### 3.4 Study Limitations and Clinical Implications

Although the results are significant, this study has several limitations. First, while the sample size met statistical requirements, it did not include a wider range of chronic inflammation-related diseases, such as rheumatoid arthritis or atherosclerosis, which limits generalizability. Second, the study mainly relied on microbiota correlation analysis and lacked direct causal validation, such as fecal microbiota transplantation or single-strain intervention, to confirm the central role of microbiota in inflammation relief. Third, lifestyle factors such as diet and physical activity, although partially controlled, may still have affected microbial abundance and immune status. The clinical implication is that GLP-1 agents may serve as a central treatment option for “metabolism–inflammation” overlap diseases, extending their use from diabetes and obesity to inflammatory liver disease, cardiovascular disease, and even autoimmune disorders. Future work should include multicenter, long-term follow-up trials to confirm their generalizability and safety across different pathological conditions. In addition, monitoring the abundance of F. prausnitzii and Roseburia spp. could serve as a microbiological indicator of treatment response, supporting the application of precision medicine.

## 4. Conclusion

This study systematically compared the effects of GLP-1 receptor agonists and NSAIDs on reducing systemic inflammation and, combined with metagenomic and metabolomic analyses, revealed potential microbiota-mediated mechanisms. The results showed that GLP-1 treatment significantly lowered inflammatory markers, including CRP, IL-6, and TNF-α, with anti-inflammatory efficacy superior to that of NSAIDs. At the same time, GLP-1 therapy markedly increased the abundance of butyrate-producing bacteria such as Faecalibacterium prausnitzii and Roseburia spp., which were negatively correlated with inflammatory indices. These findings provide direct evidence for a novel “incretin–microbiota–immune” pathway, indicating that the gut microbiota may be a key mediator of the systemic anti-inflammatory effects of GLP-1 agents. The novelty of this study lies in the first combined verification using clinical controlled experiments and microbiota analysis to establish a mechanistic link between GLP-1 therapy and microbial changes, and in proposing microbial abundance as a potential biomarker of treatment response. Unlike NSAIDs, which act mainly through the inhibition of inflammatory pathways, GLP-1 agents restore immune homeostasis through multi-level metabolic and immune regulation. This advantage provides a theoretical basis for extending their therapeutic scope to chronic inflammation-related diseases such as non-alcoholic fatty liver disease, cardiovascular disease, and immune disorders. However, this study has limitations, including a relatively small sample size, the absence of a wider disease population, and the lack of causal validation experiments. Future research should involve multicenter, long-term clinical trials, together with microbiota-targeted interventions such as probiotic supplementation or fecal microbiota transplantation, to clarify the causal role of microbiota in the anti-inflammatory effects of GLP-1 therapy. In summary, this study not only strengthens the understanding of the anti-inflammatory mechanisms of GLP-1 receptor agonists but also provides new insights for their clinical application in metabolism–inflammation overlap diseases, while highlighting the potential of microbiota abundance as a basis for precision intervention strategies.

